# Participation in a Science Festival Promotes Inclusive Science Communication around Autism Spectrum Disorder

**DOI:** 10.1101/2021.01.04.425324

**Authors:** Chris Gunter, Cynthia B Sinha, David Jaquess

## Abstract

As a high prevalence disorder with limited information about etiology, autism spectrum disorder (ASD) has been marked by confusion and miscommunication around its causes and treatments. To promote high-quality science communication, we participated in a local science festival, both providing information about the brain and ASD and asking passersby questions about their knowledge of ASD. We then asked the booth staffers to evaluate the program and conducted qualitative analyses of public and staffer responses. Public responses to the question “what would you like to know about autism?” most often concerned how the disorder was diagnosed or defined. In contrast, public responses to the question “what would you like others to know about autism?” centered around educating those unaffected by ASD on how to improve interactions and awareness, mentioning inclusivity and intersectionality, and dispelling negative stereotypes. The staffers overwhelmingly reported that in future years, they would include even more science and allow for more in-depth conversations with interested parties, as well as bringing materials in other languages. These responses are in keeping with a trend for more inclusive science communication, particularly in the field of behavioral health and ASD, and a desire to challenge myths around the condition. We conclude that our science festival interactions brought multiple benefits to public and staff.

## Introduction

Autism spectrum disorder, as defined in the Diagnostic and Statistical Manual of Mental Disorder, Fifth Edition is a neurodevelopmental disorder with key features reflecting impairment in social interactions (e.g., reciprocal communication and social interaction) and difficulties arising from overly circumscribed or repetitive interests, activities and/or behaviors (1). These disorders are present from very early in life, and they range in a spectrum of severity from mild (reflected in persistent difficulties with perspective taking and comprehending affective components of human relations) to very severe (complete withdrawal from social interaction and lack of awareness of socially mediated features of the environment). The recent epidemiological estimate of prevalence from the Centers for Disease Control and Prevention (CDC) are 1 in 54 children overall, with the rate being four times higher in males than in females (2). Age of diagnosis is crucial to receiving early treatment and affecting the trajectory of the condition; given the behavioral nature of ASD, symptoms do change over the lifespan and as a result some percentage of children may move above and below diagnostic thresholds over time (3).

In studying the apparent physical underpinnings of ASD, the scientific community has struggled to provide a complete explanation of etiology. Although some estimates point to genetics to account for up to 80% of phenotypic variance (4), the full developmental dynamic for ASD remains to be discerned. This combination of a high prevalence disorder with limited information about etiology creates a dynamic in which misinformation about etiology and treatment of the condition is rampant. While searches for the word “autism” on the internet yield hundreds of millions of hits, the quality of information varies widely and can influence, for example, the opinions of parents on accepting a diagnosis (5) or health policy makers setting standards for coverage of autism treatment (6). Healthcare consumers (as well as providers) often struggle to sort through the overwhelming volume of information to land on an empirically-informed vantage point (6,7). In sum, these are relatively common disorders for which we have limited scientific understanding of etiology, but for which there are effective assessments and empirically supported interventions that offer promising outcomes.

Public awareness, understanding, and discussion of ASD has changed substantially in the last 20 years, in part due to the sharp rise in number of diagnoses (8) and in part due to the rise in pseudoscience such as rumored links between vaccines and ASD (9). Given the 1 in 58 reported national prevalence for children meeting diagnostic criteria (10), and the increasing depiction of ASD in the media, it is very likely that most people in the US know someone or know of someone with ASD. Self-advocates have played an important role in changing the conversation as well, by describing their experiences and advocating for a shift from “autism awareness” to “autism acceptance,” for example (11). An emerging view in the advocate community is to shift from labeling ASD a “disorder” and instead to view it as part of a spectrum or even a circular rainbow of diverse brain functioning (12,13). This is in keeping with drives for inclusion and recognition of diversity in other communities.

Research suggests that these disorders are present across demographic characteristics; however, they are detected at differing rates across gender, race ethnicity, and SES. In the US there are differences in the age of diagnosis and access to care, related to race, ethnicity and socioeconomic status, as well as intersectional concerns (10,14,15). Further, people with developmental disorders like ASD tend to have reduced access to healthcare and health information, as do ethnic and racial minorities and those from lower income households (16). Thus, individuals on the autism spectrum who also come from an underserved demographic group have been particularly poorly served in the healthcare system. Given the emerging evidence base demonstrating the importance of early diagnosis and intervention in ASD (17), there is a clear need for clear and accurate communication around ASD, consistently available to everyone from specialists to the families at their moment of expressing concerns about their child’s development.

This additional element in the landscape of ASD research and communication is a dynamic of racial disparity in access to care and related delayed diagnosis and intervention is consistent with the concept of *intersectionality* in diversity—differences within differences that make a difference (18). Within any movement for equality and social justice, it is not uncommon for there to be blind spots to the differences within a recognized group. As a result, individuals whose identities fall at the intersection of classes seem to encounter doubly disparate access to care. To address these intersections, networks of self-advocates living on the autism spectrum have created alliances with disability-rights groups, LGBTQ alliances, and other movements for social justice and representation (19).

As the view of ASD has changed over time, so has the need for providers and scientists to communicate findings and recommendations around ASD. For example, erroneous reports of vaccine-autism links have taken up significant energy and sowed confusion around causes of ASD, although sites such as Vaccines Today are now providing high quality online communication and vaccine information (20). Smaller qualitative studies (21) have described a journey of progressive engagement with science for the parents of children diagnosed with ASD, whether to seek out explanations to help them understand the condition or to seek appropriate/better services for their children. A larger study in the US (22) comparing beliefs and understanding of parents and scientists regarding ASD reported that there was significant discordance between parents and scientists on their beliefs about causes of ASD and research priorities. In partial contrast, a large European study (23) found that the autism community surveyed was generally supportive of autism research, recommending that community perspectives be continually surveyed and considered in the design of research studies, including the use of preferred terminology such as “at-risk” infants. We believe that successful communication with families seeking information about ASD must include interactive and inclusive messages and activities, welcoming multiple perspectives and tailoring itself to formal or informal settings, and therefore designed our public engagement activity accordingly. This may involve intentional inclusion of words or other documentation of views from families that include a person with ASD, such as families from racial or ethnic backgrounds other than the dominant culture (e.g., Latinx, African-American, recent immigrant, east-Asian, Indian); families including parents and caregivers of many family structures; and people living on the autism spectrum of differing genders, ages, and/or races. Luisi, Rodgers and Schultz (24) argued that science communication requires training opportunities, which should include experiential learning, and they pointed out that this type of training begs a framework for program evaluation.

Given our desire to communicate high-quality scientific information about ASD, we included participation in multiple public events in the Dissemination and Outreach Core for our NIH-Funded Autism Center of Excellence grant (NIMH 2P50 MH100029). Specifically, we sought to engage with members of the public at a public science event, the Atlanta Science Festival, which closes with an “Exploration Expo” attracting up to 30,000 people. Science festivals are becoming a more common method for interaction with the public, including in areas of controversial science (25), and can be a valuable tool for evaluating public engagement with scientific topics. Our booth, entitled “How Does Your Brain Talk?”, offered both scientific information and a chance for passersby to answer specific questions about possible knowledge gaps regarding ASD. The booth also included a visual exemplar of brain development in the form of an animated portrayal of brain development research findings. Such exemplars have been found to increase the impact of science communications about autism to the general public (26). The booth was staffed by science trainees and junior faculty members who had been communicating with the public for months in the context of tours of Marcus Autism Center and were provided a one-hour seminar on how to approach the public, how to anticipate and respond to “hot topics” that might reflect misinformation, and how to communicate key pieces of information pertinent to the booth. To assess the experiential training aspect of working in the booth, we asked questions of the individuals staffing the booth about their experience and its anticipated impact on future science communication activities. As noted by Patton (27), a qualitative analysis using a theme analysis is a common first step in program evaluation. We used this type of analysis to evaluate the feedback from both groups of stakeholders: the attendees and the booth staffers.

## Statement of Ethical Review

The authors reviewed the purpose, design, execution and use of this project and determined that it fit the definition of Program Evaluation as stipulated by Emory University (http://www.irb.emory.edu/forms/review/programeval.html). The Emory University IRB’s Non-Human Subjects Research Determination Electronic Form agreed that this project was exempt from review by the university’s Institutional Review Board.

## Methods

### Terminology

In the larger community related to autism spectrum disorders, we see the terms autism and ASD both used frequently. We also see “people with autism/ASD” and “autistic people” used and would generally defer to the choice of the person involved. In this article, we will use all four of these terms.

### Booth setup

Our booth at the Atlanta Science Festival in March 2019 included a mock magnetic resonance imaging (MRI) scanner, constructed from cardboard and a table, with sound effects for how an MRI might sound through headphones. As part of the booth experience, attendees were able to lay on the table and have the staffers, primarily students or faculty from the Marcus Autism Center, move cardboard parts of the mock MRI above them while playing the sound effects and explaining how an MRI takes pictures of the brain. [No research or evaluation were conducted or collected on this completely voluntary part of the festival booth.] Afterwards (or separately if they chose not to try the mock scanner), they would be shown a rotating brain image on a monitor, depicting the wiring of circuits in the brain as found through neuroimaging studies at our Center. At the front of the booth were flip charts as described below.

### Data Collection

Data were collected to evaluate two aspects of the community outreach event. The first question of what type of information would fit the needs of the audience at the event was evaluated by asking two questions.

Question 1: “What would you like to know about autism?”

Question 2: “What would you like others to know about autism?”

These questions were posted atop two 20-inch by 23-inch flip charts in the display booth and on the wall of the booth. Staffers directed attendees’ attention to the questions as they began interacting with the display. Two corresponding methods were used to gather responses for both questions. One consisted of the staffers in the booth transcribing oral responses from attendees onto the flip charts where the questions were posted or asking the attendees to write their own response onto the flip chart. The second method was to provide small self-adhesive note pads on which attendees wrote their responses and posted them on the wall of the booth. We did not collect any information about attendees and cannot match them to their comments. Following the event, responses gather in both methods were transcribed into a single text file.

The second research question was how the staffers experienced the role of communicating with the public and what (if anything) they learned. In order to gather this information, staffers were asked to complete a questionnaire of four open-ended questions.

1. What was the main thing you learned from this experience?
2. How did you change what you were communicating over the course of the day?
3. What did you learn about people’s perception about autism or science in general?
4. What would you change about our efforts for next time?

The questionnaire was distributed via email eight days after the event, and one follow-up email prompt was sent to those who had not responded after seven days. We did this anonymously and coding was not able to match respondent to response.

### Data Analysis

Responses from the event were analyzed using content analysis approach. This analytic approach facilitated the conceptual organization of the attendee responses (28). Content analysis utilizes deductive or inductive coding. The former coding technique purposefully codes for established set of words, phrases, or concepts (29). Inductive coding develops concepts from the responses without establishing codes a priori (29), and therefore was used for this evaluation. Once conceptual coding was completed, patterns were identified, and the concepts were grouped into meaningful categories. Frequently, content analysis involves quantifying coding categories and may retain only the most prevalent (29). For this evaluation, all responses were retained for the final analysis. The lead coder (CBS) developed the coding concepts and initial categories. For inter-coder reliability, a second and third coder reviewed coding concepts and edited organization of the categories for clarity. Consensus was reached for the final analysis. Given our small data set, we did not believe utilizing a statistical measure for reliability was appropriate; our aim was 100% agreement.

As an additional means of analyzing themes in the responses from attendees, we analyzed the text of the public responses using the Semantic Word Clouds Visualization tool (30,31). We produced two separate visualizations: first (Figures 1A and 2A), we applied a simple layout of sorting by rank. Number of words was set at 50. Similarity was determined by the cosine coefficient, ranking was by term frequency, sizing was 4:3, and color was black. Second (Figures 1B and 2B), we used the layout settings for the seam carving algorithm, which determines an image based on semantic relationships and then minimizes the number of empty spaces between groups of words (31). Number of words was set at 100. Similarity was also determined by the cosine coefficient, ranking was by term frequency, sizing was 4:3, and color was ColorBrewer 2. In all figures, stop words and numbers were removed, similar words were grouped, and the shortest word was set at three letters.

**Figure 1:**
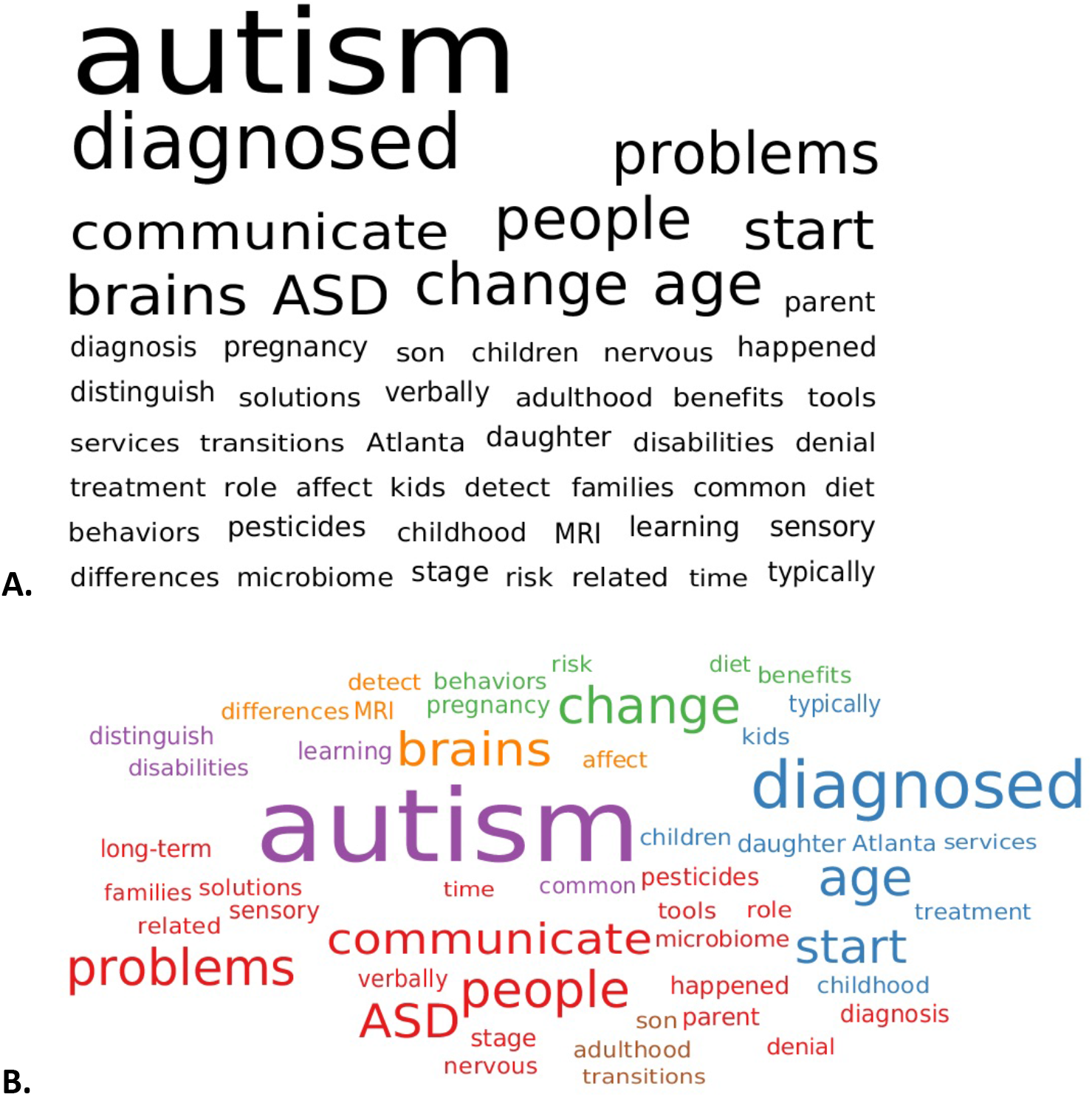
Textual analysis of answers to the question, “What do you want to know about autism?” **A:** Answers sorted by rank and term frequency. **B.** Answers sorted by seam carving algorithm and term frequency.

**Figure 2:**
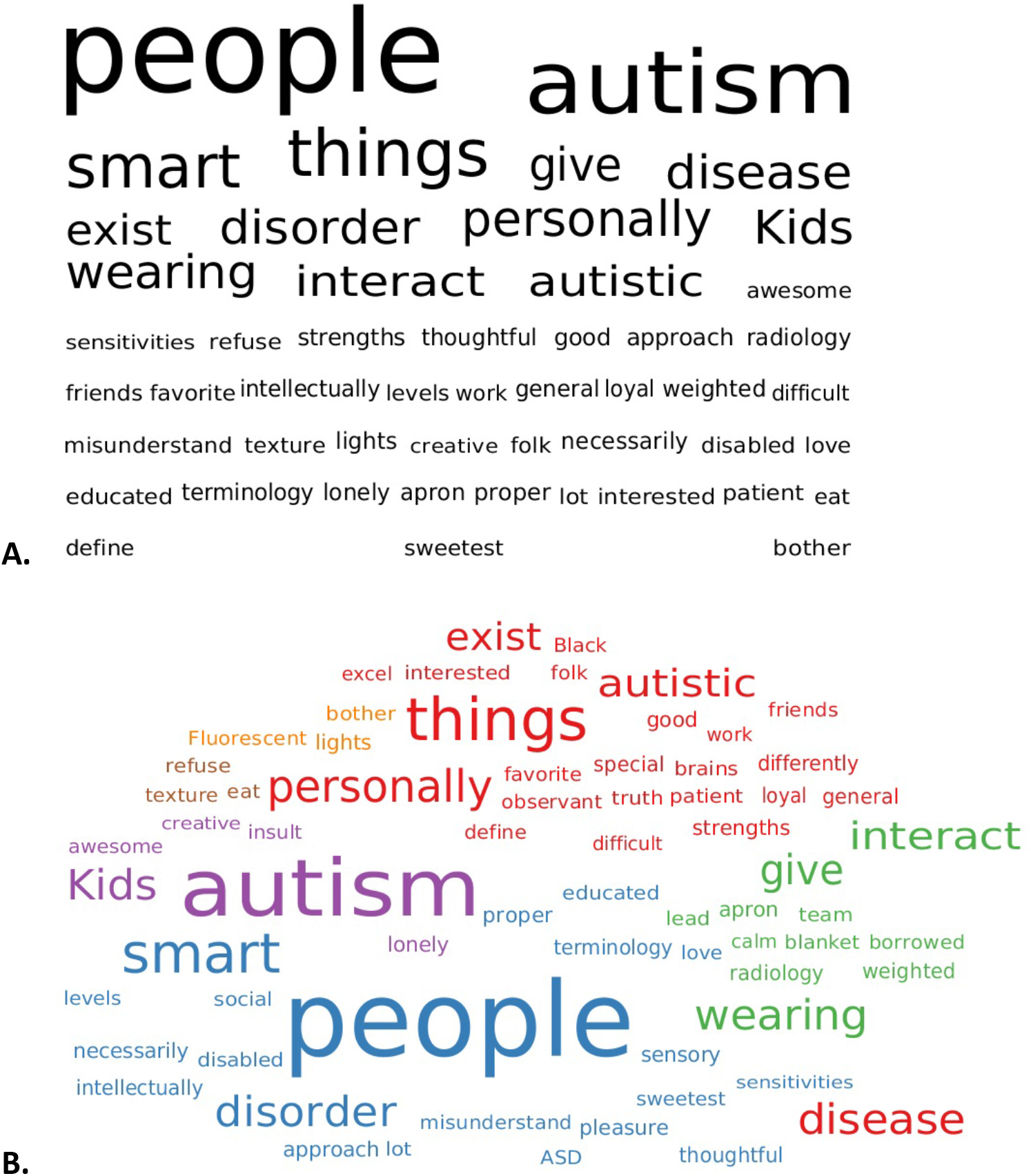
Textual analysis of answers to the question, “What do you want others to know about autism?” **A:** Answers sorted by rank and term frequency. **B.** Answers sorted by seam carving algorithm and term frequency.

## Results

Approximately 150 attendees engaged with the booth during the event; 27 responses were collected for Question 1 and 30 were collected for Question 2. The median length of response was eight and nine words for Question 1 and Question 2 respectively. Attendees appeared to include diversity in ages, gender, ethnic origin, and relationship to ASD; in discussion with the booth staffers, attendees reported themselves to be from multiple professions and family situations. Many passersby saw the name of the booth sponsor (Marcus Autism Center) and stopped by specifically to tell us about their family member, student, patient, or friend with ASD. A few individuals specifically self-identified as having ASD. Therefore, it is clear that from the general audience who self-selected by attending a science festival, our evaluation participants represent a subset with a much higher likelihood of interest in or personal relationship to ASD science.

Inductive content analysis of the attendees’ responses to our questions revealed several prominent themes. Overall, for Question 1, people appeared to be most interested in learning how ASD is diagnosed or defined. There were also multiple questions about causes for ASD, including environmental factors (but not including vaccines). Finally, specific treatment questions were raised, including service locations in the area, transitioning from childhood to adulthood and beyond, and diet or counseling requests.

For Question 2, the responses centered around educating those unaffected by ASD on how to improve interactions and awareness and dispelling negative stereotypes. Specific advice on how to treat them in the emergency room, for example, was mixed in with general advice such as “They want to interact but it’s harder for them!” Many of the statements mentioned positive characteristics like creativity, intelligence, and even “the most sweet and thoughtful people you will have the pleasure of knowing.” To combat negative labels, attendees mentioned specifically “it’s <u>not</u> a disease” and “autism does not define a person.” We also saw comments that seemed to reflect an intersectionality of marginalized experiences, e.g. “Black autistic folk exist too,” and multiple reminders of inclusivity, e.g. “It is not a disease, just a different way of thinking.” These responses appeared to have been provided by attendees who were on the autism spectrum or those that had experienced autism in their families, offering support for a subjective experience of intersectional barriers to societal inclusivity or to access of medical care.

The graphic (word cloud) analysis of responses from program attendees two the two informational questions suggested additional thematic trends. Figures 1 and 2 represent these responses in word clouds, using different analysis methods.

Figures 1A and 2A depict the top 50 words in rank order, with size of the word corresponding to its frequency in the answers. Figures 1B and 2B depict the top 100 words in a semantic word cloud, placing related words near each other and sizing each word based on its frequency in the answers. For Question 1 about what attendees would like to know, the common themes primarily centered around defining and describing the disorder (diagnosis, brain, problems, nervous, detect, problems, start) with a clear thread of words humanizing the condition (children, kids, people, age, older [individuals]; Figure 1). With regards to Question 2, the common themes include issues of increased awareness qualities of individuals with autism (many, look, smart, different, kids, Black), challenges related to the condition (sensitivities, eat, wear, trying, interact), and areas of dialogue on how to characterize the condition (disease, disorder, different; Figure 2).

All seven of the staffers responded with extensive information on the program evaluation questionnaire, with a median response length of 63 words per question, overall. The comments from the staffers reflected some of those from the attendees, including motivation to learn more about ASD themselves, and to bring more awareness about ASD to the community. A number expressed surprise at the intense level of interest in the science around ASD (and the enthusiasm of children for trying out the mock MRI scanner), but confirmed that as scientists, they enjoyed translating their scientific work and learning about the concerns and questions of those interested in ASD but not working in a scientific setting. They reported changing their approach for science communication based on the individual, sometimes focusing just on the brain and other times more on ASD.

The staffers also made specific suggestions about the booth layout: more space was needed because it became crowded at peak times, and a more sophisticated MRI scanner would be optimal. There was some disagreement at the Festival about whether it is best to reach out to people walking by instead of waiting for them to approach the booth, but in their responses, multiple staffers said the next booth should feature more engaging activities or visuals, and the team should plan to engage all passersby, perhaps with a short script. In keeping with an inclusive approach, one staffer specifically asked for some items in Spanish to be created for the next time, because “I had a few families come up to me who only spoke Spanish. I was able to translate for them, but they wanted materials to take home with them and I would’ve loved to have provided them with materials in Spanish.” Clearly, the prominent desire for next year was to include even more science and allow for more in-depth conversations with interested parties.

## Discussion

From the responses to our Question 1, “What would you like to know about autism,” the responses most often included questions on how ASD is diagnosed or defined. This result is likely to be affected by the theme of the booth, as it is related to a larger project to detect and diagnose ASD at earlier ages, and therefore this theme was much more likely to be mentioned by the staffers interacting with those entering the booth and interacting with them. In addition, we saw multiple questions about the specifics of ASD symptoms and manifestations, whether behavioral, cognitive, or social. These themes are likely influenced by the personal relationships many of our general-public attendees disclosed, including as a teacher or nurse, for example. Family members of people with autism were more likely to ask specific treatment queries, such as “Where can older children get services in Atlanta, my daughter was diagnosed at age 13.”

Overall, the responses we received when we asked what people wanted to know about ASD reflect larger questions in the community as a whole, based on our experiences interacting with the community for multiple years as a researcher (CG) and clinician (DJ). How can we help children as they transition into adulthood? What does scientific research tell us about ASD symptoms, diagnoses, and treatments? What are the specific symptoms of ASD, and what do terms like “high-functioning” mean? And what is new research, such as into the microbiome or the role of pesticides, telling us about ASD?

We did not record any questions about vaccines, and none were written on the flip-chart boards or Post-It notes. This suggests that either attendees asked those questions directly to booth staff (and would have been told that there is no link between vaccines and autism), and/or that the public who stopped by did not have this question or did not want to put it into writing.

For Question 2, on what others should know about autism, we saw many more references to inclusivity and the more personal side of the condition. Overwhelmingly, respondents want to teach those who are unaffected on appropriate ways to interact with individuals with ASD. This included advice on patience and not “taking things personally,” as well as specifics like “think twice before wearing cologne/perfume” due to sensory sensitivities. Respondents wanted to share the positive attributes of people they knew with ASD and counter the idea that autism and intellectual disability are conflated (four comments specifically mentioned that people with autism are “smart” or that they “aren’t necessarily intellectually disabled!”).

Individuality of people with ASD was a prominent theme as well, echoing themes we have heard from self-advocates for years like “Not all kids with autism look the same.” Simultaneously, there was also a theme of request for inclusion, such as “We exist too,” “We’re not that different,” and “Many people with autism are lonely….Can be good friends!”

Finally, we heard about the intersectionality of ASD with the experience of being a minority, from comments like “Black autistic folk exist too.” We grouped this into a theme with requests for “seeing” the autistic person, including other statements like “Just because they don’t ‘look’ autistic doesn’t mean they aren’t.”

We conclude from our content analyses that the science festival brought multiple benefits to our efforts to communicate with the public. At the same time, there is clearly value to the ASD community in seeing themselves represented at such an event. Some self-identified autistic individuals stayed for in-depth conversations and shared their experiences with us. Our staffers reported they enjoyed learning from those with ASD too, and that they would use the experience to shape future research and future interactions. In addition, their responses did seem to reflect growth in science communication through expanded knowledge of public need and increased repertoire for engaging with them. The greater variety of hands-on activities in the booth suggested by staffers could function as multiple visual exemplars and thus support increased learning for attendees (26). In future festival booths, we would also like to recruit some staffers or volunteers who identify as being autistic themselves.

The primary limitation of this work is the difficulty in generalization from our findings, given its nature as an evaluation of a one-time event. This evaluation may also provide guidance for future research in this area. Data collection could be expanded either by repeating the same event as we had planned to do (unfortunately, the 2020 Atlanta Science Festival was canceled due to the coronavirus pandemic) or repeating this setup at another festival to increase the sample size. These added events would allow a more refined analysis of responses and possible exploration of issues related to intersectionality. Researchers might need to gather data from multiple events or in multiple locations within a science fair to achieve this goal. Based on the themes derived from the present evaluation, future research could formalize a set of information to be communicated to meet the needs identified here; such studies should include a measure of efficacy in getting the message across to members of the public. It also could be informative to include different methods of communication (perhaps including different language translations) within a single event in order to compare the efficacy of those methods. In addition, designing a study that gathered contact information would allow researchers to follow-up and gather data about the durability of information communicated to attendees. Each of these directions could further expand our understanding of effective science communication.

## Author Contributions

CG and DJ designed the evaluation and the questionnaires. CS completed the first round of coding, and all authors participated in later iterations of analysis. All authors contributed to manuscript revision, read and approved the submitted version.

## Acknowledgments

We thank all of the attendees at our Atlanta Science Festival exhibit, the Festival organizers, and the volunteers at the exhibit: Aiden Ford, Skai Glasser, Emma Goldman, Cheryl Klaiman, Longchuan Li, Sarah Markert, Adriana Mendez, Jack Olmstead, Tristan Ponzo, and Sarah Shultz. CG also thanks the Fondation Brocher.

